# OriGene: A Self-Evolving Virtual Disease Biologist Automating Therapeutic Target Discovery

**DOI:** 10.1101/2025.06.03.657658

**Authors:** Zhongyue Zhang, Zijie Qiu, Yingcheng Wu, Shuya Li, Dingyan Wang, Yong Liu, Zhuomin Zhou, Yusong Hu, Yuhan Chen, Duo An, Yongbo Wang, Yu Li, Zhenyi Zhong, Chubin Ou, Zichen Wang, Feng Tang, Jack Xiaoyu Chen, Runmin Ma, Jun Li, Xinyu Wang, Wei Lu, HaoMing Xue, Wenxin Zhang, Zhijian Wei, Runze Ma, Zhanpeng Shi, Ke Wang, Qingwu Liu, Bo Dong, Yuexi He, Tinghai Liu, Jiayu Gu, Shuotong Song, Qiantai Feng, Jian Zhang, Bo Zhang, Luyi Tian, Lei Bai, Qiang Gao, Siqi Sun, Shuangjia Zheng

## Abstract

Here, we present OriGene, a self-evolving multi-agent system that functions as a virtual disease biologist, systematically identifying original and mechanistically grounded therapeutic targets at scale. OriGene’s architecture integrates over 600 specialized tools through a Model Context Protocol (MCP), enabling it to reason across diverse data modalities including genomics, protein networks, pharmacology, clinical records and literature evidence, to generate and prioritize target discovery hypotheses. We implemented a strategy combining a knowledge graph-based Tool RAG with an advanced agent selection mechanism to enable dynamic, context-aware tool deployment. Through a self-evolving framework, OriGene continuously integrates human and experimental feedback to iteratively refine its core thinking templates, tool composition, and analytical protocols, thereby enhancing both accuracy and adaptability over time. To comprehensively evaluate its performance, we established TRQA, an original benchmark comprising over 1,900 expert-level question-answer pairs spanning a wide range of diseases and target classes. OriGene consistently outperforms human experts, leading research agents, and state-of-the-art large language models in accuracy, recall, and robustness, particularly under conditions of data sparsity or noise. Critically, OriGene nominated previously underexplored therapeutic targets for liver (GPR160) and colorectal cancer (ARG2), which demonstrated significant anti-tumor activity in patient-derived organoid and tumor fragment models mirroring human clinical exposures. These findings demonstrate OriGene’s potential as a scalable and adaptive platform for AI-driven discovery of mechanistically grounded therapeutic targets, offering a new paradigm to accelerate drug development.

## 1. Introduction

The identification of therapeutic targets is a central step in modern drug discovery, enabling tailored interventions for complex diseases. Despite decades of technological progress, over 90% of drug candidates fail during clinical development[47, 38]. In many cases, these failures arise not from issues with the compounds themselves, but from flawed initial hypotheses about the biological role, disease relevance, or druggability of the chosen targets[11, 15], and the target-selection process is still fragmented and heavily reliant on intuition. Critical data and analytical tools for key modalities, including genomics, transcriptomics, proteomics, metabolomics, imaging, and clinical records, generally operate in isolation[2, 40, 9], limiting cross-modal reasoning and mechanistic integration. Their outputs are often incompatible, and few frameworks enable unified interpretation across domains. This fragmentation results in a slow, costly, and failure-prone path from molecular insight to viable therapeutic hypothesis. The substantial cost of experimental validation, frequently exceeding $2 million per target for methods like CRISPR screens or organoid assays[24, 32], further constrains the number of hypotheses that can be rigorously tested, worsening a critical bottleneck in early-stage drug development.

Recent advances in large language models (LLMs) have introduced agent-based frameworks capable of automating scientific reasoning [18, 31, 21, 5], coordinating diverse tool [45, 44, 13, 34], and integrating knowledge from literature with structured data [30, 12, 18, 16]. While these systems have laid important groundwork, they are not purpose-built for the unique complexities of therapeutic target discovery and validation. This process demands structured, cross-scale reasoning that connects molecular mechanisms with disease pathology, pharmacological modulation, and competitive therapeutic landscape [20, 14]. Expert disease biologists typically engage in iterative cycles of literature mining, mechanistic inference, tool-driven analysis, and experimental refinement to generate and evaluate hypotheses[12, 42]. An ideal AI agent to support this process must integrate heterogeneous data, interpret results within a deep biological context, and dynamically adapt its reasoning in response to both human expert and experimental feedback. Current AI systems offer limited evidence of possessing this full suite of capabilities with the necessary depth and rigor, and the absence of standardized benchmarks curated for target discovery further hinders their systematic evaluation and development.

To address these gaps, we present OriGene, a self-evolving multi-agent system designed to function as a virtual disease biologist, focusing on the systematic identification of original and mechanistically grounded therapeutic targets. OriGene integrates over 600 expert tools and curated biomedical databases within a unified framework that supports multi-modal reasoning across genomics, transcriptomics, proteomics, phenomics, and pharmacological data. It autonomously generates and refines therapeutic hypotheses by analyzing functional interaction networks, disease ontologies, perturbation data, and literature-derived evidence. A core feature of OriGene is its capacity to self-improve over time. It refines its responses to individual queries through iterative reasoning cycles and progressively enhances its broader analytical capabilities by learning from accumulated experience and experimental feedback.

We evaluated OriGene against expert biologists, leading general-purpose language models, and existing specialized AI agents using the TRQA, a custom-curated benchmark of 1,921 question-answer pairs covering diverse disease areas and target classes. OriGene consistently outperformed all baselines in key metrics such as accuracy, recall, and robustness, particularly when faced with data sparsity or noise. As proof-of-concept for its discovery capabilities, OriGene identified and prioritized previously underexplored therapeutic targets, GPR160 for hepatocellular carcinoma and ARG2 for colorectal cancer. These agent-driven nominations were subsequently validated experimentally using patient-derived organoids, tumor fragment models, and relevant in vivo models, confirming activity analogous to that observed in clinical settings. Together, these results demonstrate that OriGene’s self-evolving, multi-agent architecture provides a robust foundation for scalable, mechanistic, and clinically relevant therapeutic target discovery.

## 2. Results

### 2.1. OriGene automates therapeutic target discovery via self-evolving multi-agent reasoning

OriGene is a self-evolving multi-agent system that serves as a virtual disease biologist, designed to identify original and mechanistically grounded therapeutic targets at scale. As shown in Fig.**1**A, OriGene integrates over 600 expert tools and curated biomedical databases across fundamental biology, disease biology, pharmacology, and competitive landscape. Its self-evolving framework orchestrates specialized agents—Coordinator, Planning, Reasoning, Critic, and Reporting, within a tightly coupled workflow. These agents transform complex biological queries into structured hypotheses and actionable insights, ensuring transparent reasoning for expert review and collaboration.

**Fig. 1:**
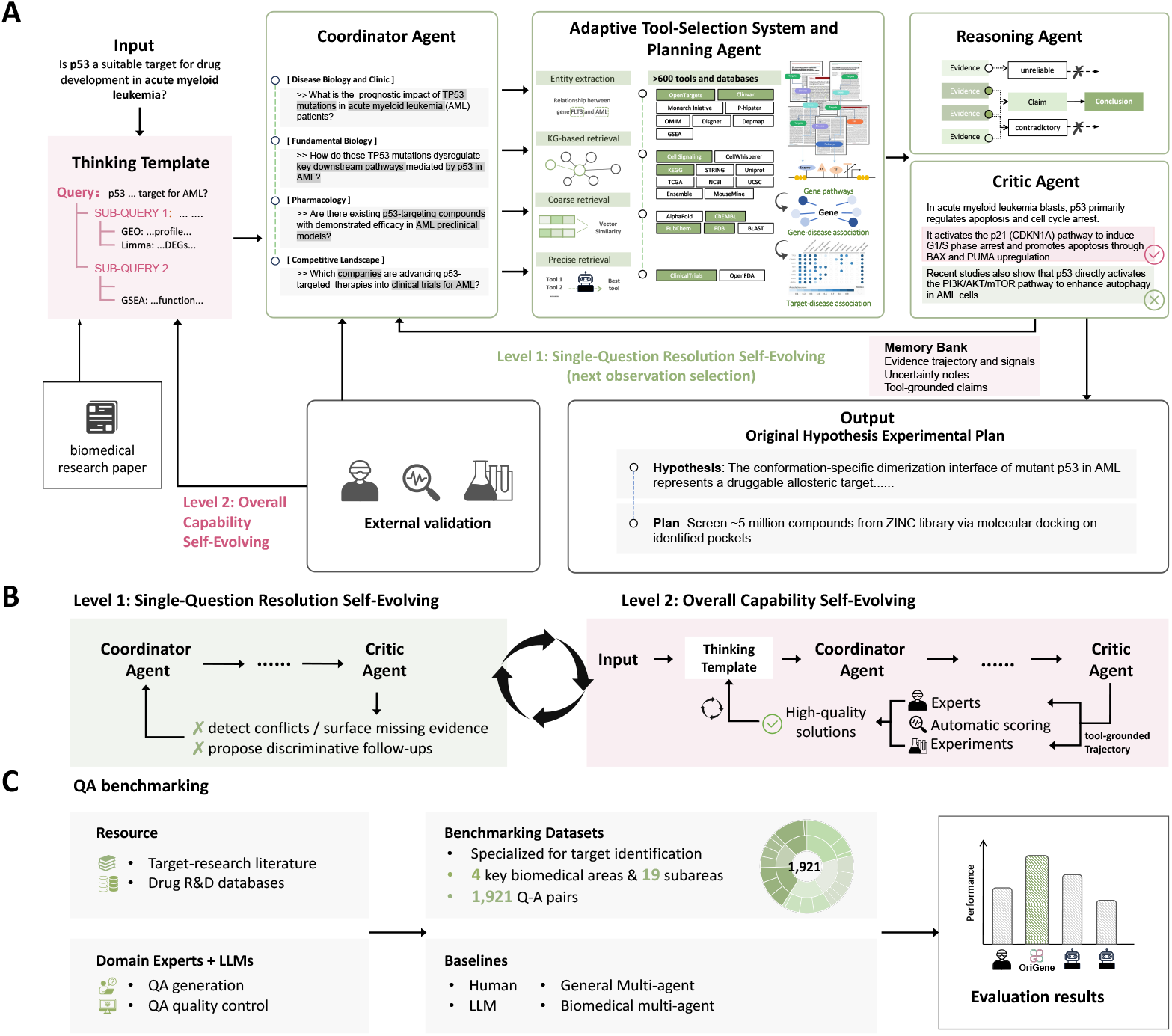
The overall framework, self-evolving mechanism, and benchmarking process of OriGene. **(A)** OriGene deploys interconnected agents (Coordinator, Planning, Reasoning, Critic) to generate original hypotheses, decompose subqueries, select and utilize tools. These outputs undergo iterative refinement through dynamic interaction with experts and experimental systems. **(B)** OriGene exhibits self-evolving capabilities on two levels: iteratively refining answers to complex queries and expanding system-wide expertise through multigenerational template evolution. **(C)** Evaluation schemes, including QA benchmark datasets assessing broad aspects of target identification capabilities.

Upon receiving a target research question, the **Coordinator Agent** analyzes and systematically decomposes it into focused sub-problems, guided by domain-specific **thinking templates**—structured logical frameworks distilled from target research literature that ensure scientific rigor and adherence to best practices in problem breakdown. For each sub-problem, the **Adaptive Tool-Selection System and Planning Agent** autonomously determines which specialized biomedical databases, computational tools, or analytical strategies to invoke, dynamically sequencing and integrating resources to flexibly address the unique requirements of each sub-query. The **Reasoning Agent** then synthesizes and compresses multi-modal outputs, identifying not only key relationships but also potential contradictions among gene targets, diseases, molecules, and signaling pathways. The **Critic Agent** performs rigorous gap analysis, evaluating both the completeness and scientific soundness of the emerging solution, and provides targeted feedback to drive further refinement. Throughout this process, a centralized memory bank accumulates and organizes both raw data and processed evidence, supporting transparency and robust knowledge management within the workflow. Leveraging the accumulated evidence and multi-round reasoning outputs, the **Report Agent** compiles and articulates the final output as a coherent and structured scientific report. This workflow was implemented on a mixed-model backbone, with GPT-41 driving the Coordinator, Planning, Critic, and Reporting agents, and GPT-41-mini managing low-latency post-processing of tool outputs. More architecture details can be found in Method 4.1.

A defining feature of OriGene is its **self-evolving capability** on two levels, as illustrated in Fig.**1**B. First, during the resolution of each query, OriGene engages in iterative cycles of task decomposition, tool utilization, reasoning, and reflection—enabling test-time scaling, where additional computational resources and more iterations directly translate to improved response quality. Second, OriGene continuously enhances its system-wide reasoning ability through the expansion and refinement of its template library: it systematically extracts new, high-quality thinking templates from its own most effective solution trajectories, annotated by human experts or through experimental validation, facilitating a virtuous cycle of capability growth. Details of self-evolution are provided in Method 4.3.

This recursive, template-guided, and self-evolving multi-agent architecture empowers OriGene to systematically address complex biomedical questions, autonomously select and employ diverse disease biology tools, and continuously advance its own scientific reasoning abilities through cumulative experience.

### 2.2. OriGene outperforms human and other AI models in target research reasoning

To assess OriGene’s capabilities in therapeutic target discovery and assessment, we first evaluated its performance on the Target Research Question-Answering (TRQA) benchmark, which we constructed to test advanced reasoning and information retrieval in this domain (Fig.**2**A-B, Method 4.4). TRQA consists of two subsets: TRQA-lit, which focuses on biological research from literature and includes 172 multiple-choice and 1,108 short-answer questions covering fundamental biology, disease biology, clinical medicine, and pharmacology (Fig.**3**A, left); and TRQA-db, which centers on the competitive landscape with 641 short-answer questions regarding drug R&D pipelines and clinical trials (Fig.**3**A, right). Together, these datasets span a wide range of targets and diseases (Fig.**3**C), enabling systematic evaluation. Detailed construction procedures for TRQA are described in Methods 4.4. We employed distinct evaluation strategies for multiple-choice and short-answer questions, also detailed in Methods 4.4.

**Fig. 2:**
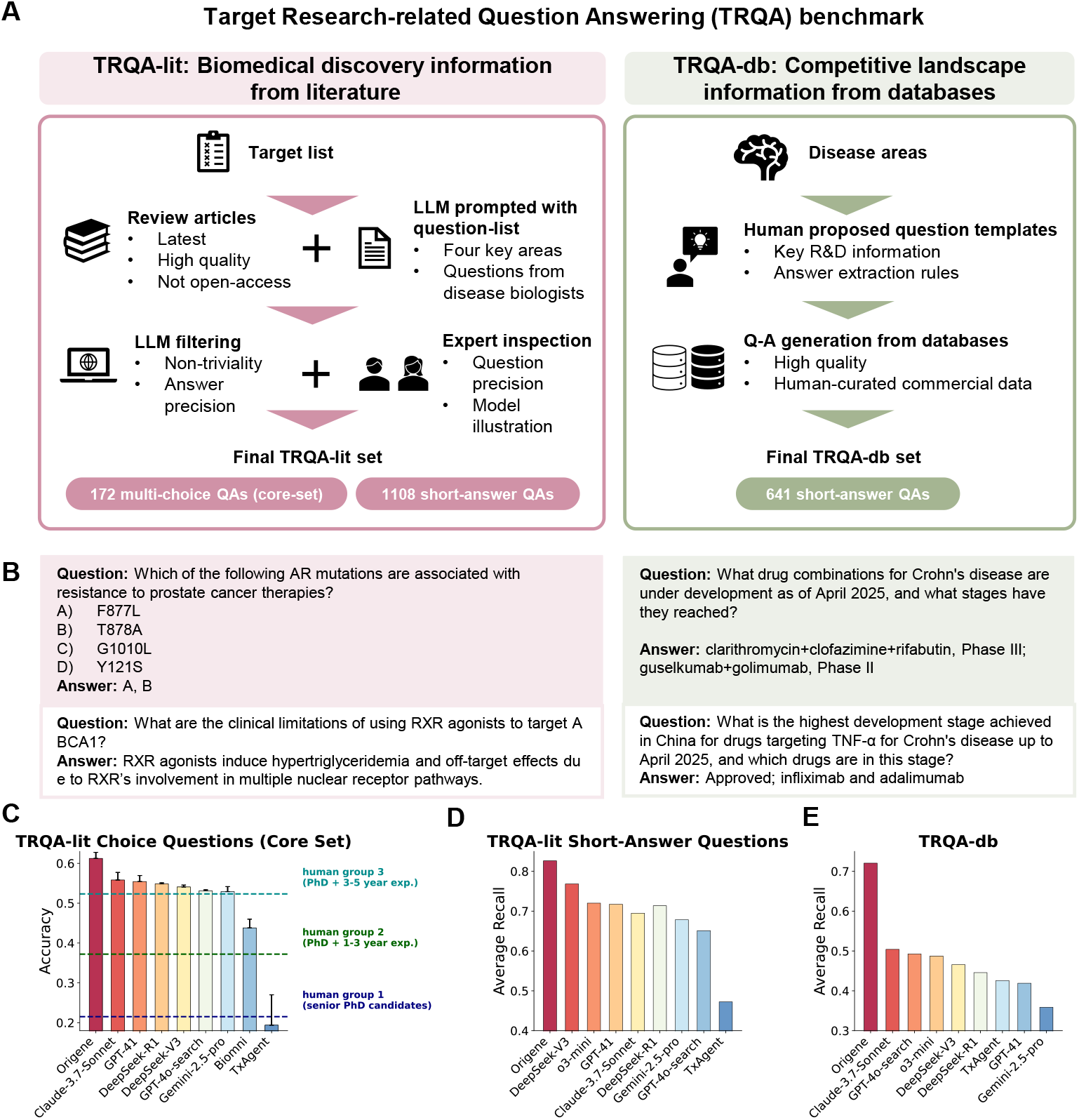
Target Research-related Question Answering (TRQA) benchmark for evaluating biomedical knowledge and target identification skillsets. **(A)** Overview of TRQA benchmark comprising two complementary datasets: TRQA-lit derives questions from biomedical literature through LLM filtering and expert curation, yielding 172 multiple-choice and 1,108 short-answer questions; TRQA-db generates 641 short-answer questions from structured databases using human-designed templates and extraction rules. **(B)** Representative question examples from both datasets with corresponding answers. **(C-E)** Performance evaluation results showing accuracy for multiple-choice questions **(C)** and average recall for short-answer questions from literature **(D)** and database sources **(E)** across different systems, with human performance benchmarks indicated for reference groups of varying expertise levels.

**Fig. 3:**
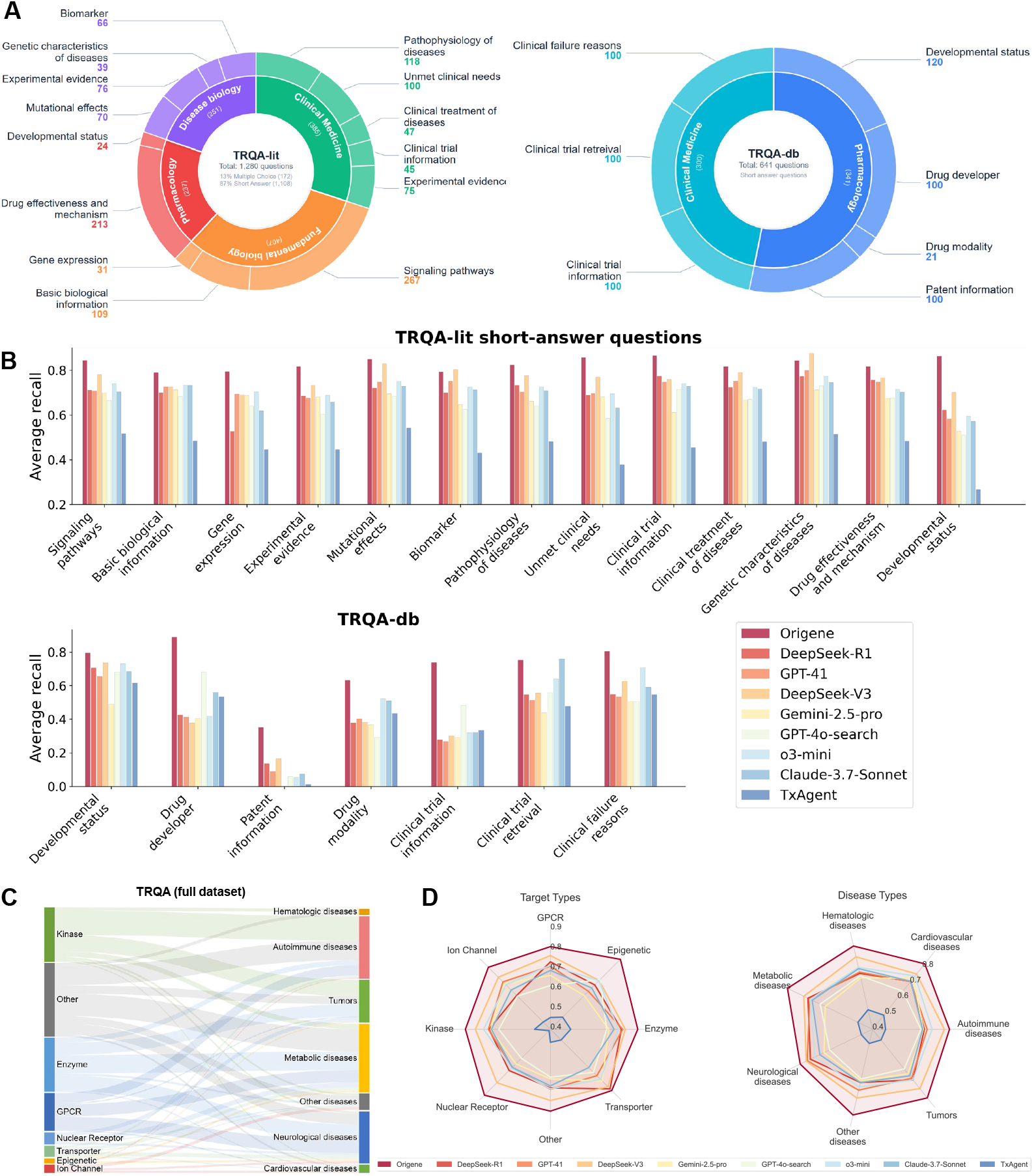
Domain-stratified performance analysis on TRQA benchmark. **(A)** Question distribution across biomedical categories for TRQA-lit (left, n=1,280) and TRQA-db (right, n=641). **(B)** Average recall performance stratified by biomedical domains, demonstrating domain-specific competencies of OriGene relative to baseline systems. **(C)** Associations between drug target classifications and disease categories across the dataset. **(D)** Average recall across target types (left) and disease types (right).

As shown in Fig.**3**, on the TRQA-lit (multiple choice) benchmark, OriGene achieved a score of 60.65. This performance is particularly noteworthy as it surpassed the scores achieved by human domain experts, who were grouped by experience level and permitted to use Google search (as detailed in Methods and Fig.**2**C). OriGene also outperformed leading AI competitors on this task, including DeepSeek-V3[29], DeepSeek-R1[19], and GPT-41[23], while substantially exceeding the specialized TxAgent[13]. We also evaluated the concurrently released agent, Biomni[22]. Given the timing, this analysis was confined to the TRQA-lit benchmark, where a rapid evaluation on a core set reaffirmed OriGene’s significant performance advantage. OriGene continued its strong performance on the TRQA-lit (short answer) benchmark, designed to assess the generation of concise and accurate literature-derived responses, scoring 82.76%, which again surpassed all other models, including DeepSeek-V3 and GPT-41. Furthermore, in navigating complex relational data for competitive landscape analysis, OriGene achieved the highest score of 72.05% on the TRQA-db benchmark. This significantly outperformed models like Claude-3.7-sonnet[3] and GPT-4o-search, underscoring its robust capabilities in extracting and reasoning over structured, domain-specific database information. OriGene’s consistent strengths across different biomedical domains within TRQA are further detailed in Fig.**3**B.

To contextualize these findings against broader AI reasoning capabilities, we further validated OriGene’s performance on several public benchmarks: GPQA[39], DbQA[27], and LitQA[27]. On GPQA, which requires complex reasoning and integration of external information to answer graduate-level scientific questions, OriGene demonstrated strong performance, scoring 78.20% overall. This surpassed Gemini-2.5-pro[17], a cluster of models including DeepSeek-R1, DeepSeek-V3, and o3-mini[37], as well as GPT-41 and GPT-4o-search, highlighting OriGene’s effective utilization of external information. The DbQA benchmark, testing specific database querying or structured data reasoning, showed OriGene as the clear top performer with the highest score of 50.00%. This placed it significantly ahead of models like DeepSeek-R1 and o3-mini, Claude-3.7-Sonnet, Gemini-2.5-pro, and DeepSeek-V3, highlighting OriGene’s notable advantage in structured data interaction and tool integration. Consistent with its strong literature-based performance on TRQA-lit, OriGene again achieved the highest score on the LitQA benchmark, which assesses biomedical literature comprehension. This performance substantially outpaced other leading models such as DeepSeek-R1, o3-mini, GPT-41, and Gemini-2.5-pro, further demonstrating its advanced ability to process and reason over knowledge-intensive textual data critical for scientific applications.

Overall, OriGene’s performance on both our custom TRQA benchmark, where it exceeded human expert benchmarks on relevant tasks, and established public datasets confirms its advanced capabilities in biomedical reasoning, information synthesis from structured and unstructured text, and robust handling of domain-specific knowledge, often surpassing state-of-the-art LLMs and specialized research agents.

### 2.3. OriGene’s performance improves with computation and tool access

OriGene’s architecture, featuring an iterative loop between Coordinator and Critic Agents, is designed for flexible scaling of test-time computation across two key dimensions: processing depth, through more iterations of refinement, and width, by increasing the number of sub-queries explored concurrently. This adaptability is vital for addressing the varying complexity of biological research questions. Our empirical study confirms that increasing this test-time computation consistently enhances OriGene’s benchmark performance. This effect was clearly demonstrated on the LitQA dataset, which assesses literature-based question answering. As illustrated in Fig.**4**B (left panel), OriGene’s accuracy rose steadily from 62.81% when using minimal computation (defined as “1x”, representing a single sub-query) to 78.39% when the computational budget was increased ninefold (“9x”). While the performance gains began to plateau around the “9x” mark, indicating a point of diminishing returns for this specific task, these results clearly show OriGene’s capacity to leverage additional computational resources at inference time to deepen its analysis and improve response quality.

**Fig. 4:**
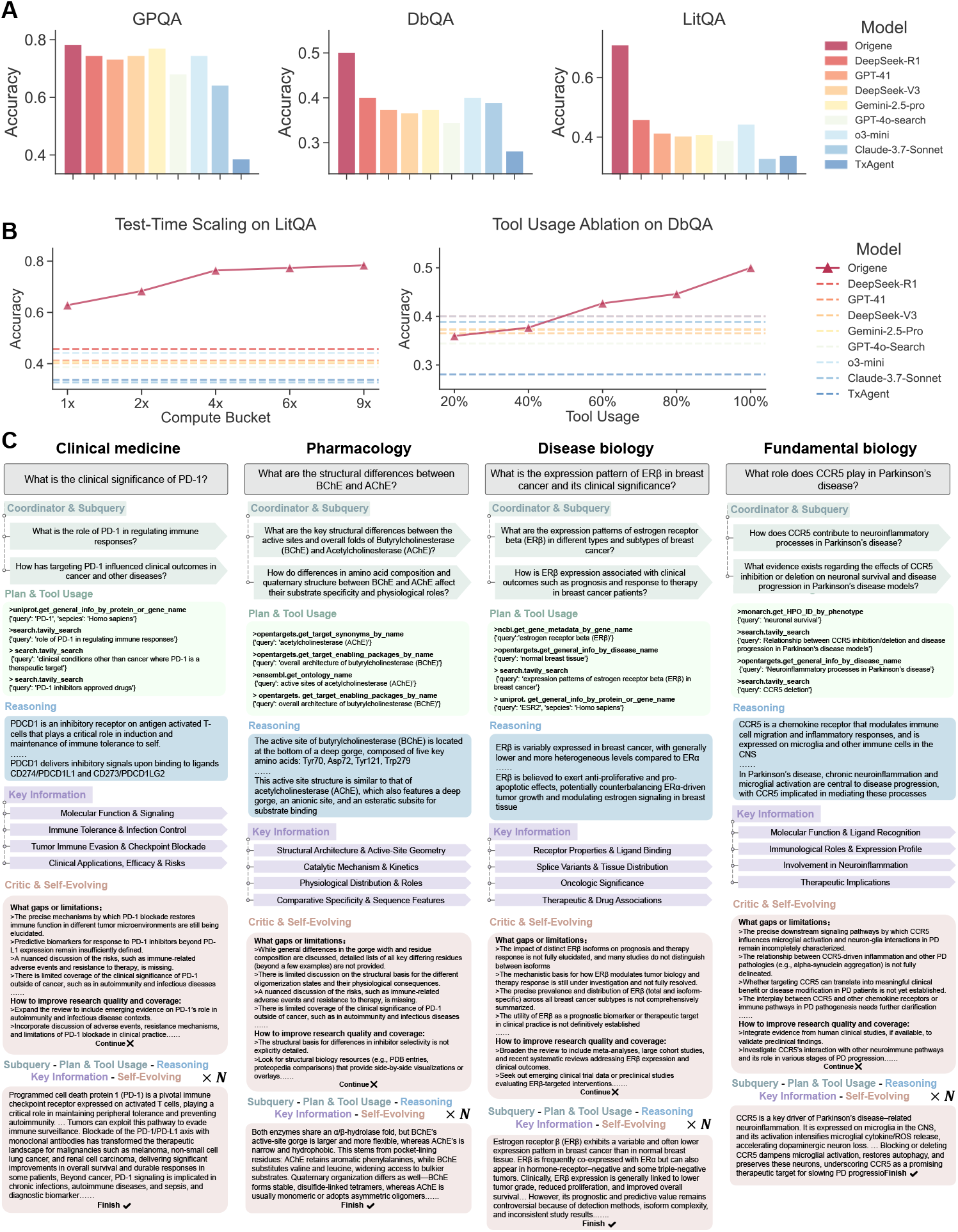
Comparative evaluation and ablation analysis of biomedical QA models across multiple benchmarks. **(A)** Accuracy performance of OriGene and baseline large language models on three QA tasks: GPQA (left), DbQA (middle), and LitQA (right). **(B)** Left: Test-time scaling effects on LitQA, compared to static baseline performances. Right: Tool usage ablation on DbQA with baselines shown for reference. **(C)** Self-Evolving OriGene framework for systematic biological enquiry. Detailed breakdown of the four sub-agent components: “Clinical significance of PD-1,” “Structural comparison & pharmacokinetics,” “Pediatric use,” and “Target modulation & pathway inhibition.” For each track, the figure summarizes planning and tool selection, sub-query execution, evidence analysis, key insights, and subsequent self-evolution steps.

Given OriGene’s integration of numerous specialized biomedical tools (detailed in Methods 4.2), we also conducted an ablation study to precisely quantify the impact of tool availability on its problem-solving capabilities. To achieve this, we systematically varied tool access by randomly disabling a predetermined percentage of OriGene’s available tools for each question presented from the DbQA dataset, ensuring that different tools were disabled across questions to generalize the findings. This study revealed a strong and direct correlation between the comprehensiveness of the available toolset and OriGene’s overall problem-solving effectiveness. Specifically, as depicted in Fig.**4**B (right panel), OriGene’s accuracy on the DbQA benchmark, which evaluates database querying and structured data reasoning, declined substantially from 50.00% when all tools were accessible to 35.96% when its access was restricted to only 20% of the tools. This underscores how critical integrated, disease-specific tools are for real-world target reasoning, whereas generalist agents lack this modular robustness.

To demonstrate how OriGene orchestrates external tools for open-ended biomedical questions, we highlight four representative ‘agent trajectories’ spanning diverse research domains (Fig.**4**C). Each trajectory initiates with a natural-language query, for instance, regarding the clinical relevance of PD-1 or the structural divergence between cholinesterases. OriGene’s capabilities are showcased across various applications. It integrates disease-gene data with pathway analysis to connect PD-1 dysfunction to inflammatory processes in clinical medicine, explains enzyme substrate specificity by analyzing structural data in pharmacology, identifies potential biomarkers by synthesizing expression profiling and gene-disease associations in disease biology, and elucidates signaling pathways by correlating diverse data sources in fundamental biology.

Throughout these applications, OriGene’s ‘self-evolving’ mechanism identifies knowledge gaps, such as limited transcriptomic coverage for certain ESR2 variants, and automatically initiates follow-up queries using relevant databases or analytical modules. These results highlight the system’s capacity to process heterogeneous biomedical data, maintain evidence provenance, and iteratively refine its outputs without human intervention, contributing to its observed performance.

Together, these findings highlight critical operational characteristics of OriGene: its problem-solving capacity is not static but can be significantly amplified by greater computational investment during inference, and its effectiveness is critically dependent on comprehensive access to its integrated suite of specialized biomedical tools.

### 2.4. OriGene’s self-evolving discovery of GPR160 as a novel target in hepatocellular carcinoma

Beyond benchmark validation, we deployed OriGene to the real-world task of identifying novel therapeutic targets within complex human malignancies. OriGene’s capacity to autonomously interrogate and synthesize vast biomedical datasets—spanning literature, clinical records, and multi-omics databases—was leveraged to nominate previously uncharacterized candidate targets in distinct solid tumors.

In the context of hepatocellular carcinoma (HCC), a malignancy with significant unmet medical needs, OriGene was tasked with uncovering novel, druggable target candidates. Through an iterative, non-human dominant agent-driven process encompassing information retrieval from diverse resources (including PubMed[46], GEPIA[28], UCSC[35], TCGA[43], GEO[6], and OpenTarget[8]), multi-modal data analysis, and mechanistic hypothesis generation (Fig.**5**A), OriGene initiated its reasoning to identify therapeutically relevant targets. This comprehensive bioinformatic pipeline enabled OriGene to prioritize G-protein coupled receptor 160 (GPR160) from an extensive pool of 125 initial candidates with multiple rounds of prompt iterations (Fig.**5**A-B). Subsequent agent-led interrogation of large-scale clinical datasets revealed marked GPR160 upregulation in HCC tumor tissues relative to minimal expression in adjacent non-malignant liver (Fig.**5**B). Critically, OriGene’s analysis of CPTAC data established a significant inverse correlation between elevated GPR160 expression and recurrence-free survival (RFS) in HCC patients (p=0.0057, Fig.**5**C), underscoring its clinical relevance and OriGene’s ability to autonomously triage multiple targets to clinical relevant candidates.

**Fig. 5:**
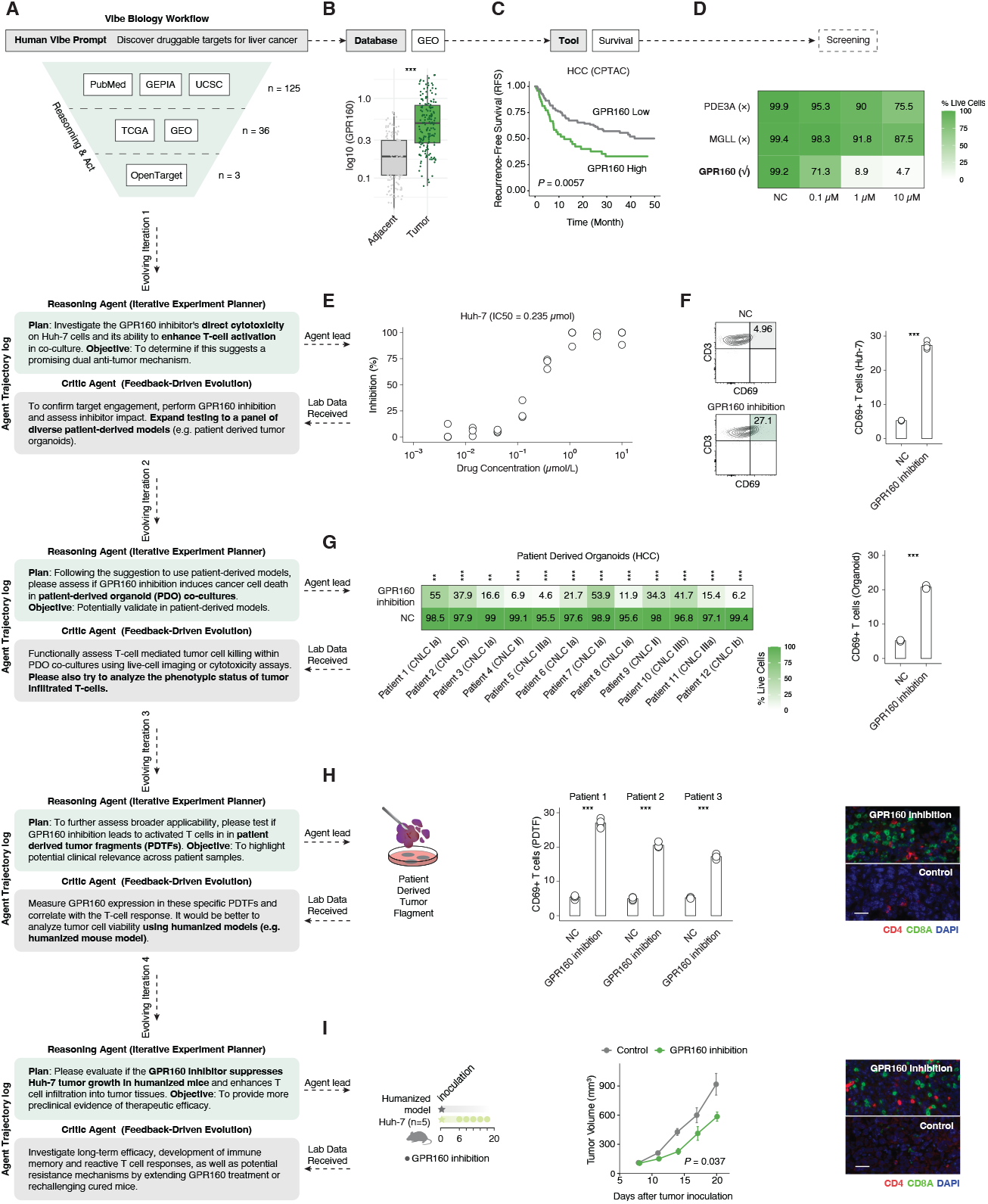
OriGene-driven discovery and validation of GPR160 as a therapeutic target in HCC. **(A)** OriGene’s initial prompt and data sources for target discovery in HCC. Iterative reasoning and action cycles involving the Reasoning Agent and Critic Agent guide the discovery process. **(B)** GPR160 mRNA expression (log10 scaled) in HCC tumor versus adjacent normal tissues from GEO datasets. **(C)** Kaplan-Meier analysis of recurrence-free survival (RFS) in HCC patients from the CPTAC cohort, stratified by GPR160 expression. **(D)** Cell viability screening results showing the effect of GPR160 inhibitor compared to negative controls (NC) and controls including PDE3A, MGLL. **(E)** Dose-response curve of GPR160 inhibition in Huh-7 HCC cells (n = 3), yielding an IC50 of 0.235 µmol/L. **(F)** Representative flow cytometry plots (CD69 expression on CD3+ T cells) and quantification of CD69+ T cells in Huh-7 co-cultures treated with or without GPR160 inhibitor (n = 4). **(G)** Left: Effect of GPR160 inhibition on cell viability in a panel of HCC patient-derived organoids (PDOs) (Patients 1-12, representing different CNLC stages). Right: Quantification of CD69+ T cells in PDO co-cultures treated with or without GPR160 inhibitor. **(H)** GPR160 inhibition promotes T-cell activation in patient-derived tumor fragments (PDTFs). **(I)** GPR160 inhibition suppresses tumor growth and enhances T-cell infiltration in a humanized mouse model (Scale bar: 50 µm).

Following the autonomous, agent-driven identification of GPR160, OriGene formulated an experimental plan to investigate the direct cytotoxicity of GPR160 inhibition and its capacity to modulate T-cell activation. An undisclosed deep learning–enabled screening campaign was conducted, leading to the rapid identification of a candidate small-molecule inhibitor within four weeks. Subsequent pharmacological characterization in vitro, guided by OriGene’s experimental design, demonstrated that GPR160 inhibition induced potent effects in Huh-7 cells (Fig.**5**D-E) as well as in the HepG2 HCC cell line. To assess efficacy in models more reflective of in vivo tumor heterogeneity, in response to OriGene’s iterative feedback loop, which suggested evaluation in patient-derived systems for enhanced clinical relevance, GPR160 inhibition was examined across a panel of patient-derived organoids (PDOs) (Fig.**5**G). Employing co-culture systems of cancer cells with T-lymphocytes, GPR160 inhibition was observed to promote marked T-cell activation (Fig.**5**F-G). Progressing to an even more complex *ex vivo* system, treatment of patient-derived tumor fragments (PDTFs) with the GPR160 inhibitor similarly resulted in an increased proportion of activated CD69+ T cells across multiple patient samples, with representative immunofluorescence imaging indicating enhanced CD4+ and CD8+ T-cell infiltration (Fig.**5**H).

Finally, to evaluate the *in vivo* efficacy and the dual mechanism of GPR160 inhibition, OriGene proposed experiments using a humanized mouse model bearing Huh-7. Systemic administration of the GPR160 inhibitor led to a significant suppression of tumor growth compared to the control group (p=0.037 at day 20, Fig.**5**I, middle panel). Crucially, immunofluorescence analysis of tumor sections from these mice revealed a marked increase in the infiltration of both CD4+ and CD8+ T-cells into the tumor microenvironment of GPR160 inhibitor-treated animals (Fig.**5**I, right panel), with no significant change in mouse weight during treatment. Taken together, these data generated through OriGene’s iterative reasoning and automated experimental planning establish GPR160 as a compelling therapeutic target in HCC, whose inhibition mediates direct tumor cell cytotoxicity while concurrently fostering potent anti-tumor immune responses.

### 2.5. Vibe Biology-enabled, self-evolving prioritization by OriGene validates ARG2 as a therapeutic target in colorectal cancer

Extending this automated discovery paradigm, OriGene was tasked with identifying therapeutic targets in colorectal cancer (CRC) to orchestrate a multi-stage, self-evolving experimental validation strategy (Fig.**6**A). The system’s initial bioinformatic analyses, drawing from multiple MCPs such as PubMed, GEPIA, UCSC, TCGA, and OpenTarget, converged to prioritize Arginase-2 (ARG2) from 86 candidates as the lead candidate from a larger pool of possibilities. OriGene’s reasoning highlighted significant ARG2 overexpression in comprehensive CRC patient cohorts compared to adjacent normal tissues (Fig.**6**B). Supporting proteomic evidence, including data from the Human Protein Atlas (e.g., CAB009435), confirmed ARG2 protein upregulation in CRC tissues (Fig.**6**C).

**Fig. 6:**
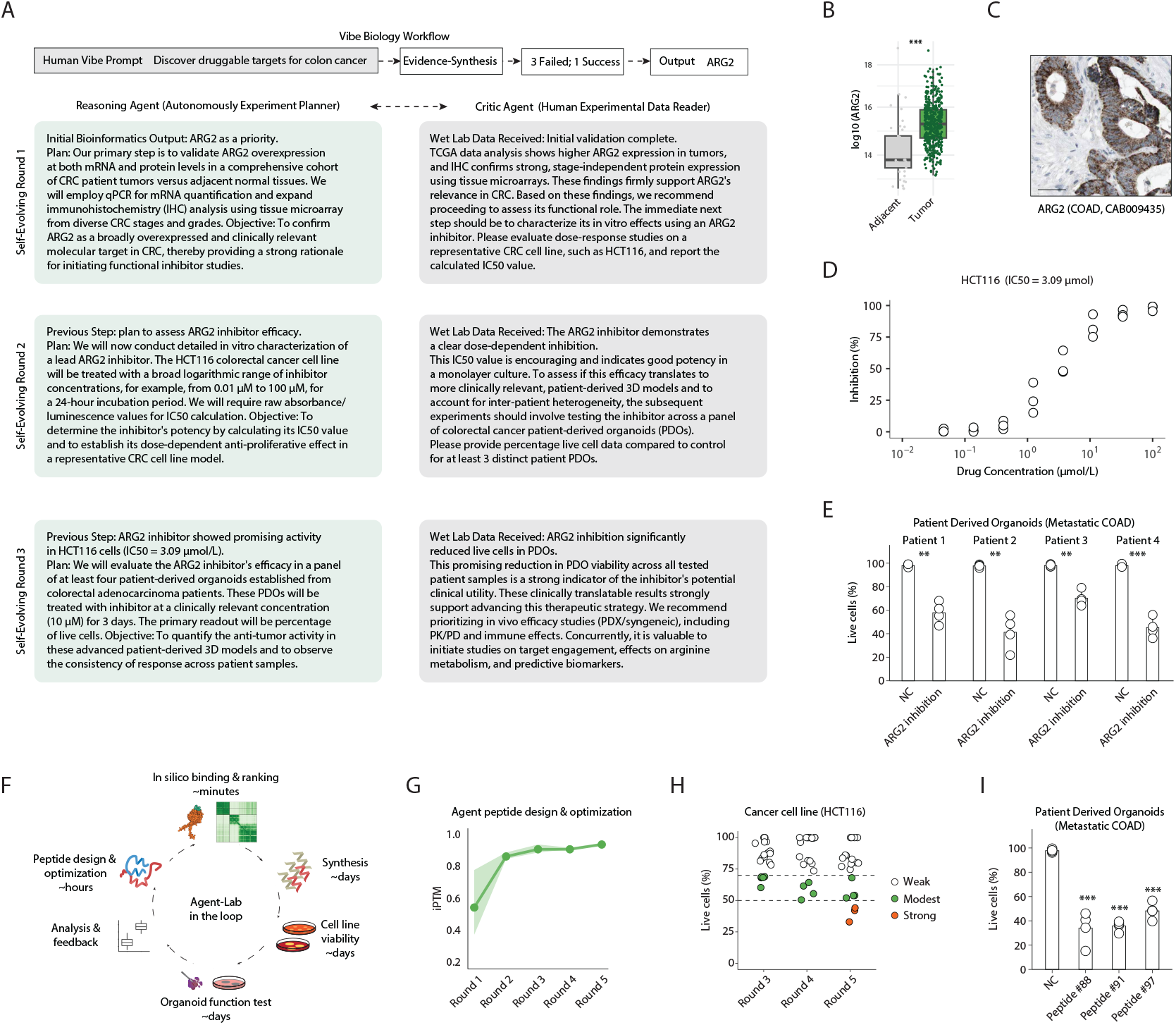
OriGene’s autonomous discovery and initial validation of ARG2 as a therapeutic target in CRC. **(A)** Schematic of OriGene’s iterative, multi-round process for CRC target discovery, involving initial bioinformatics analysis, Reasoning Agent and Critic Agent (Experimental Data Reader) interactions. The prompt directs discovery for colorectal cancer, querying databases and outputting ARG2 after filtering (3 failed, 1 success shown illustratively). **(B)** ARG2 expression in CRC tumor versus adjacent normal tissues from the TCGA dataset (**p *<* 0.001; Wilcox test). **(C)** Representative immunohistochemistry (IHC) staining of ARG2 in human colorectal adenocarcinoma (COAD) tissue (Human Protein Atlas, CAB009435). **(D)** Dose-response curve of ARG2 inhibition in HCT116 CRC cells, demonstrating an IC50 of 3.09 µmol/L. This step is part of OriGene’s “Self-Evolving Round 2” experimental plan. **(E)** Effect of ARG2 inhibition (10 µM) on cell viability in a panel of four patient-derived organoids (PDOs) from metastatic COAD. Data presented as percentage live cells relative to no-drug control (NC) for each patient (Patient 1-4). This validation is part of OriGene’s “Self-Evolving Round 3”. Error bars represent mean ± SD. **(F)** Schematic of the AI-driven, closed-loop discovery cycle for peptide optimization, integrating in silico design, chemical synthesis, and functional testing in organoids. **(G)** Agent-led multi-round optimization of peptide inhibitors, showing improvement in the iPTM score over five rounds. **(H)** Inhibitory effects on the HCT116 cancer cell line during multi-round peptide optimization. The plot shows the effects of different peptides on cell viability across rounds 3, 4, and 5, categorized by their inhibitory effect as Weak, Modest, or Strong. **(I)** Optimized lead peptides demonstrate significant anti-tumor activity in patient-derived organoids from metastatic colorectal adenocarcinoma (metastatic COAD). Compared to the negative control (NC), peptides #88, #91, and #97 all significantly reduce the proportion of live cells in the organoids. Data are presented as mean ± SD, n=3. ***p < 0.005.

Based on this robust, agent-driven prioritization, OriGene formulated a strategy to evaluate the efficacy of ARG2 inhibition. Subsequent experimental data confirmed that a selective ARG2 inhibitor elicited a clear dose-dependent inhibition of HCT116 CRC cell proliferation in multiple cell lines (Fig.**6**D). Demonstrating its adaptive approach, OriGene’s self-evolving experimental protocol (as depicted by “Self-Evolving Round 2” leading to “Self-Evolving Round 3” in Fig.**6**A, D, E) then mandated validation in more clinically relevant patient-derived models. Consequently, the ARG2 inhibitor was assessed in a panel of four patient-derived organoids (PDOs) established from metastatic colorectal adenocarcinoma. The inhibitor demonstrated significant anti-tumor activity, reducing viable cell populations across these distinct PDOs (Fig.**6**E). This iterative cycle, wherein OriGene’s autonomous reasoning guided experimental design and subsequent data interpretation, successfully identified and provided initial preclinical validation for ARG2 as a novel therapeutic target in CRC.

We next leveraged the platform to guide therapeutic development through an ‘Vibe Biology-enabled Agent-Lab in the loop’ optimization of ARG2-targeting peptides (Fig.**6**F), where humans provide only monitoring while AI autonomously iterates design and validation. This iterative process integrated agent-led in silico design with wet-lab validation. After initial computational screening (Rounds 1–2; Fig.**6**G), experimental data from each round informed the subsequent redesign. For instance, following viability assays in HCT116 cells in Round 3, the agent identified and fixed a key structural motif from the top-performing peptides. In Round 4, it then re-optimized flanking regions to improve affinity. This iterative refinement culminated in three highly potent peptides in Round 5, which strongly inhibited HCT116 cell viability (Fig.**6**H). Crucially, these lead candidates (Peptides #88, #91 and #97) also demonstrated robust anti-tumour efficacy in metastatic COAD patient-derived organoids (Fig.**6**I), validating the framework’s ability to autonomously drive the discovery of potent, clinically relevant molecules.

In summary, OriGene’s application to distinct solid tumors, exemplified by HCC and CRC, not only uncovered GPR160 and ARG2 as promising, druggable targets but also powerfully demonstrated the robustness and transformative potential of our AI-driven discovery engine. These findings underscore OriGene’s capability as a self-evolving virtual disease biologist to substantially accelerate future therapeutic development by automating key stages of target discovery.

## 3. Discussion

In this study, we introduce OriGene, a self-evolving multi-agent system representing a novel paradigm for therapeutic target discovery. Unlike general-purpose AI co-scientists, OriGene is engineered with deep domain-specific reasoning capabilities and an architecture optimized for identifying mechanistically grounded targets. To enable objective evaluation in this complicated area, we established TRQA, an open-source benchmark comprising 1,921 disease- and target-specific questions, setting a new standard for assessing such AI agents. OriGene coordinates a network of specialized agents (Coordinator, Planning, Reasoning, Critic, and Reporting) that reason over multimodal data, leveraging OriGeneHub—an integrated ecosystem of 645 expert tools and 18 curated biomedical databases. This unified framework bridges fundamental biology, disease biology, pharmacological, and clinical evidence, allowing for an interpretable and iterative process of hypothesis generation and refinement crucial for modern drug discovery.

Our results demonstrate that OriGene achieves strong performance across key stages of target discovery, consistently outperforming human experts (even when aided by search tools) and leading biomedical AI agents on benchmark tasks. This robust performance stems from its transparent and interpretable reasoning, leading to efficient and explainable automated discovery. In experimental validation, OriGene successfully identified and prioritized previously underexplored targets, GPR160 in hepatocellular carcinoma and ARG2 in colorectal cancer. These AI-driven nominations were subsequently confirmed through experimental testing in patient-derived organoid and tumor fragment models, demonstrating efficacy in humanized settings. These findings illustrate OriGene’s ability to translate vast, heterogeneous knowledge into actionable and experimentally relevant therapeutic hypotheses.

Target discovery remains an inherently complex, multi-modal challenge where current foundation models often struggle without task-specific designs like OriGene’s template-guided reasoning, tool integration, and feedback-driven adaptation. By augmenting expert scientific workflows, OriGene achieves strong benchmark performance and generates experimentally relevant hypotheses. However, further development is needed to address OriGene’s current limitations: i) its extensive tool set requires pre-defined rules and post-training techniques for improved efficiency and reduced topic drift; ii) its discovery potential could be broadened by expanding data sources including structured patents and clinically annotated datasets; and iii) its generated hypotheses necessitate refined scoring and filtering to address challenges in safety, testability, originality, and biological plausibility. The continued development of more robust foundational models and self-improving strategies will also be crucial for advancing OriGene’s capabilities.

Overall, this study represents a significant step toward systematically accelerating therapeutic target discovery using advanced, agent-based AI systems. We demonstrate that by deeply integrating domain knowledge and mimicking the complex reasoning and decision-making processes of disease biologists, OriGene can generate novel, actionable, and mechanistically grounded hypotheses across diverse disease contexts. As generative AI models and biomedical data resources continue to mature, frameworks like OriGene are poised to provide a foundational AI-first platform for scalable, end-to-end drug discovery, advancing the transformative potential of precision medicine.

## 4. Methods

### 4.1. OriGene System Architecture and Core Agents

The Virtual Disease Biologist framework is driven by a suite of interconnected agents, each designed to tackle distinct aspects of the research process with precision and rigor. As shown in Fig.**1**A, these components—Coordinator, Planning, Reasoning, Critic, and Reporting Agents—work in concert to transform complex biological questions into actionable insights for drug discovery. Information flows seamlessly between these components via a disease biology-specialized memory bank that centralizes critical evidence connecting gene targets, diseases, molecules, and signaling pathways. By integrating adaptive planning, sophisticated analysis, and stringent quality control, this system ensures a comprehensive, scientifically robust approach to target evaluation. Below, each agent’s role is detailed to illustrate how they collectively advance the research process with clarity and efficiency. Unless otherwise noted, inference used a mixed-model backbone: GPT-41 powered the Coordinator, Planning, Critic and Reporting agents, and GPT-41-mini handled low-latency post-processing of tool outputs.

#### Coordinator Agent

This central agent specializes in analyzing and breaking down user queries into logical sub-problems. It examines incoming questions, identifies key components that need investigation, and structures them into manageable sub-questions. The agent ensures that complex inquiries are systematically addressed by organizing them into a coherent analytical framework. During subsequent iterations, the Coordinator Agent reviews feedback from the Critic Agent and evaluates existing information in the Memory Bank to refine its approach. This reflective process enables it to formulate improved questions that address gaps in understanding or explore promising new directions. By continuously adapting its query decomposition strategy based on accumulated knowledge and critical feedback, the Coordinator Agent progressively enhances the depth and relevance of the investigation. This iterative refinement ensures that the questioning process evolves intelligently throughout the conversation, leading to more comprehensive and accurate responses.

#### Planning Agent

This agent focuses on developing strategic execution plans to address the sub-queries identified by the Coordinator Agent. Rather than formulating questions, the Planning Agent determines which specialized tools and resources should be deployed to effectively resolve each sub-query. In collaboration with the **Adaptive Tool-Selection System** (see Method 4.2), it selects the most appropriate tools from a pool of over 600 biomedical tools based on the contextual demands of each sub-query. It then generates executable tool-calling statements that specify how each selected tool should be used. After implementing these tool-calling strategies, the Planning Agent systematically records the acquired knowledge in the Memory Bank for future reference. This structured approach to resource utilization ensures that each sub-query is addressed with the most suitable analytical methods, creating an efficient pipeline from question decomposition to knowledge acquisition and storage.

#### Reasoning Agent

This agent serves as the analytical processor of collected information, drawing from the knowledge repository established by the Planning Agent. It performs critical comparative analysis across multiple sub-queries, identifying patterns, connections, and inconsistencies within the assembled data. The Reasoning Agent excels at synthesizing evidence by recognizing and consolidating complementary information while flagging contradictory findings for further investigation. A key function of this agent is information compression - strategically condensing extensive tool-generated outputs to reduce context overhead without sacrificing essential insights. During this compression process, the Reasoning Agent prioritizes relationships between critical entities such as targets, diseases, molecules, and signaling pathways, ensuring that these vital connections remain prominent in the streamlined knowledge representation. By transforming raw data into cohesive, context-optimized insights, the Reasoning Agent creates the foundation for comprehensive understanding while maintaining analytical efficiency throughout the investigation process. Finally, the Reasoning Agent updates the Memory Bank with these processed results, ensuring that the refined and consolidated knowledge becomes available for subsequent analytical cycles and agent interactions.

#### Critic Agent

This agent functions as an evaluative overseer, critically examining the alignment between user queries and the information accumulated in the Memory Bank. It performs a gap analysis by comparing the original questions with the current state of knowledge, identifying areas where information remains incomplete, uncertain, or insufficiently addressed. The Critic Agent methodically assesses whether the collective findings adequately resolve each aspect of the user’s inquiry, highlighting specific dimensions requiring further investigation or clarification. By pinpointing weaknesses in the current understanding, whether from contradictory evidence, limited data sources, or unexplored angles, this agent drives continuous improvement in the response quality. The Critic Agent’s reflective assessment becomes a crucial feedback mechanism that guides subsequent iterations, ensuring that knowledge gaps are systematically addressed and that the final response achieves comprehensive coverage of the user’s information needs. This critical evaluation process maintains intellectual honesty by acknowledging limitations while simultaneously directing efforts toward developing increasingly complete and nuanced answers.

#### Reporting Agent

This agent transforms complex research findings into clear, visually rich reports tailored to different audiences. It organizes information logically based on the specific target, adjusts technical content for readers ranging from scientists to executives, creates layered summaries that balance detail with accessibility, includes interactive visualizations, links conclusions to supporting evidence, and highlights knowledge gaps with specific recommendations for further research. This thoughtful approach ensures that sophisticated target assessments can be effectively communicated across teams and departments.

#### Thinking Template-Guided Reasoning

To address the challenge of logical hallucinations in disease biology reasoning, we augment the Coordinator Agent and Planning Agent with structured thinking templates. While tool integration effectively reduces factual hallucinations, the complex logical processes required for query decomposition and tool selection still rely heavily on the base model’s reasoning capabilities, which can introduce domain-specific logical errors.

Our solution implements a template-guided approach using a curated collection of thinking templates extracted from bioinformatics research papers that focus on in silico methodologies. These templates capture expert reasoning patterns by analyzing the logical architecture of published research. Using large language models, we systematically extract and formalize how researchers decompose complex biological questions into investigative components, select appropriate computational tools for each sub-query, and establish purposeful connections between analytical steps.

These thinking templates provide domain-specific scaffolding that guides the agents’ reasoning processes, ensuring that query decomposition and tool selection follow established scientific methodologies rather than relying solely on the model’s general reasoning capabilities. This approach significantly enhances the agents’ ability to emulate expert-level logical reasoning in complex biological investigations while reducing the occurrence of logical hallucinations.

#### Comprehensive Safety and Ethical Oversight

Safety and ethical responsibility are foundational to the OriGene system’s architecture. The framework implements continuous, multi-layered monitoring that spans the entire research workflow—from initial user queries through sub-query decomposition, data analysis, and reporting. Every user question is systematically screened to detect potentially malicious intent, such as attempts to generate harmful biological insights or access sensitive data in ways that could violate ethical or legal standards. As queries are broken down into sub-queries, each analytical step is further evaluated in real time to ensure it does not inadvertently promote unethical research directions, privacy violations, or misuse of biomedical resources. These checks are automatically performed using a combination of large language models and embedded ethical rule sets. When potential risks are identified—such as dual-use research of concern, unauthorized data access, or exploration of ethically sensitive topics—the process is interrupted, concerns are documented, and unsafe outputs are prevented. In addition, all findings and recommendations are delivered with transparent annotations of any safety considerations or uncertainties. Through this rigorous, proactive oversight, OriGene ensures that every aspect of the research process is conducted with scientific integrity, ethical responsibility, and regulatory compliance.

### 4.2. The OriGene Agentic Tool System

OriGene integrates 645 biomedical tools to deliver real-time insights across four key categories: disease biology, fundamental biology, pharmacology, and competitive landscape, to precisely mimic a disease biologist’s reasoning workflow. At its core, OriGene uses an advanced Agent2Agent architecture. It allows individual agents to independently engage with specific biomedical tools, processing natural language queries, formulating requests, and transforming raw data into clear, relevant answers.

#### Toolset Construction Strategy

The selection of tools and databases integrated into OriGene is driven by the core design principle of emulating the reasoning workflow of a disease biologist. We deliberately focused on incorporating capabilities within the four primary functional domains that human experts prioritize during target discovery and validation: Disease Biology, Fundamental Biology, Pharmacology, and Competitive Landscape. These domains represent the essential facets of expert evaluation. Consequently, the model utilizes specialized toolsets within each domain:

- Disease Biology Tools: To evaluate the functional impact of gene perturbations within cellular and disease contexts, this domain leverages key resources including ClinVar[26] for assessing the clinical relevance of genetic variants, GSEA (Gene Set Enrichment Analysis)[41] for interpreting perturbational signatures (such as those from CRISPR, RNAi, and LINCS) within the context of biological pathways and disease states, and other perturbational datasets. These tools collectively enable the interpretation of gene function and its disruption in disease mechanisms.
- Fundamental Biology Tools: To infer disease-relevant causal mechanisms, this domain leverages core biological knowledge resources, including UniProt[10] for comprehensive protein annotation and functional characterization, Gene Ontology (GO)[4] for standardized functional annotation, KEGG[25] for detailed pathway analysis, foundational AI models (e.g., AlphaFold3[1]) for analysis the structure of target, protein-protein interaction networks for mapping mechanistic cascades and identifying functional modules.
- Pharmacology Tools: To systematically assess target tractability and drug development potential, this domain integrates key resources including bioactivity databases (e.g., ChEMBL [33]), structural repositories (e.g., PDB[7]) and specialized computational toolkits—collectively enabling multi-dimensional therapeutic evaluation.
- Competitive Landscape Tools: To evaluate commercial viability and strategic positioning, this domain utilizes specialized resources—including patent analytics platforms, commercial intelligence databases, and pipeline tracking systems—to identify approved drugs, clinical-stage candidates, licensing deals, and partnership activities. The final toolset balances domain coverage with source diversity, ensuring that each stage of the biomedical reasoning process is supported by high-quality, task-appropriate computational resources.

#### Adaptive Tool-Selection in Agent2Agent Framework

To address the challenge of selecting from over 600 tools in our Agent2Agent framework, we developed a strategy to prevent agent distraction from an overly broad context of tool descriptions. We implemented a two-stage retrieval-augmented generation (Tool-RAG) workflow that couples a domain-specific knowledge graph with LLM reasoning. This knowledge graph-based approach ensures a focused and precise retrieval process, enabling the Coordinator Agent to identify the most appropriate tools for each sub-question.

- Knowledge-graph construction. First, we build a specialized knowledge graph. The nodes of this graph represent 23 key biological entity classes, such as *Drug, Disease* and *miRNA*. The edges connecting these nodes represent tool actions, linking their required input entities to their output entities. Crucially, both nodes and edges store a list of “related tools,” creating an efficient index that maps biological entities and their transformations to the specific tools that can act upon them. Once the graph is in place, a two-stage process selects the optimal tool for a given query.
- Stage 1: General purpose selector. The system first attempts to answer the query using broadly applicable “general” tools for common lookups related to molecules, proteins, diseases, or a domain-specific web search. The LLM translates the natural-language sub-question into the structured input required by these tools. This streamlined approach quickly handles most routine queries with minimal delay.
- Stage 2: Expert selector. For queries demanding more specialized functionality, an Expert Selector provides a necessary supplement to the General Selector. This involves a more detailed two-step retrieval process. First, a **coarse retrieval** step is performed. The remaining entities and relationships are mapped onto the knowledge graph. The union of their adjacent “related tools” sets forms a preliminary tool pool. To narrow this pool, we compute the semantic similarity between the description of each tool and the query context. This similarity is computed using embeddings generated by a separate embedding-only LLM. Second, a **precise retrieval** step follows: for each candidate tool identified during coarse retrieval, the LLM is given a prompt containing the query fragment and the tool description; then, it selects the most appropriate tool and produces the exact parameterization required for execution.

This hierarchical, knowledge-graph–guided strategy ensures that each Tool Agent delivers precise, interpretable, and contextually relevant answers while seamlessly supporting the Agent2Agent pipeline for complex biomedical reasoning and data integration.

### 4.3. Self-Evolving Mechanisms of OriGene

We demonstrate that OriGene virtual biologist agent possesses self-evolving capabilities, enabling potential performance improvements with sufficient computational resources. This self-evolution operates on two distinct levels: within single-question resolution (Fig.**1**B, left) and across overall capability development (Fig.**1**B, right). The agent iteratively refines its responses to individual queries while simultaneously enhancing its comprehensive performance through accumulated experience and knowledge integration.

#### Level 1: Single-Question Resolution Self-Evolving

At the first level, OriGene demonstrates test-time scaling through an iterative self-improvement process when addressing individual research questions. Similar to the DeepResearch approach, our agent employs a structured cycle of task decomposition, tool utilization, reflection, and re-planning to progressively enhance its response quality.

When presented with a complex biological query, the coordinator agent first decomposes it into manageable sub-problems. Then the planning agent strategically deploys appropriate tools from its toolkit to gather relevant information. Crucially, the reasoning agent and the critic agent continuously evaluate the quality and relevance of intermediate results through a reflection mechanism. This critical self-assessment enables OriGene to identify limitations in its current approach, correct misconceptions, and reformulate its solution strategy accordingly. The planning of each problem is grounded in a problem-specific cognitive template. This template is dynamically refined throughout the planning process by incorporating the user’s query and accumulated research knowledge, thereby enabling the system to self-evolve in response to the demands of a specific task.

This iterative cycle continues until convergence, with each iteration producing incrementally improved outputs. Our experiments demonstrate that providing additional computational resources for more iterations consistently yields enhanced response quality, particularly for complex biological research questions requiring deep domain knowledge. Importantly, this self-evolving capability emerges without additional training, representing a form of emergent intelligence that scales with computational investment at inference time.

#### Level 2: Overall Capability Self-Evolving

Beyond single-question improvements, OriGene exhibits a second level of self-evolution through continuous enhancement of its system-wide capabilities. Central to this process is OriGene’s thinking template retrieval mechanism, which undergoes progressive refinement through multiple generations of templates.

The initial corpus of thinking templates was extracted from published literature, providing OriGene with foundational reasoning patterns for biological research. However, the system is designed to transcend these initial capabilities through an autonomous expansion process. Each template is instantiated multiple times to sample diverse reasoning trajectories, facilitating test-time scaling via multi-trajectory interaction. As OriGene processes user queries and benchmark problems, it generates high-quality solutions through its Level 1 test-time scaling mechanism. These solutions, particularly those demonstrating novel or exceptionally effective reasoning approaches, are systematically analyzed to extract new thinking templates.

This creates a virtuous cycle of capability expansion: high-quality outputs from earlier iterations become inputs for template extraction, generating a second generation of templates with enhanced reasoning patterns. These new templates are then incorporated into the coordinator agent’s retrieval database, enabling more sophisticated approaches to subsequent problems(Algorithm.**??**). This process continues iteratively, with each generation of templates building upon insights from previous solutions.

Our experiments demonstrate that this multi-generational template evolution leads to measurable improvements in OriGene’s performance across diverse biological research tasks. Importantly, this capability development occurs without traditional model retraining, instead leveraging the system’s own outputs as a form of self-supervised learning. This approach enables OriGene to continuously adapt to emerging research methodologies and domain knowledge, representing a significant advancement toward truly autonomous scientific agents.

### 4.4. Benchmark Design, Baselines, and Evaluation Methods

#### Construction Details of TRQA Dataset

To assess OriGene’s comprehensive capacities, which involve addressing specific tasks across key areas of therapeutic target discovery—including fundamental biology, disease biology, pharmacology, and clinical medicine—it is crucial to evaluate its ability to conduct effective planning, gather useful information, choose appropriate tools, reason to scientific conclusions, and critically self-evolve. This evaluation must consider information from both extensive research literature and competitive landscape data related to drug R&D pipelines and clinical trials. Recognizing that existing biomedical benchmarks often focus on general knowledge and may lack the specific depth required for therapeutic target discovery, we established the Target Research Question-Answering (TRQA) benchmark to bridge this gap and systematically evaluate OriGene alongside other multi-agent frameworks. The TRQA dataset consists of two subsets: TRQA-lit, a literature-based QA set focusing on the biological research of therapeutic targets, and TRQA-db, a database-derived QA set prioritizing the competitive landscape relevant to targets of interest. The construction of both datasets involved a comprehensive panel of human experts and LLM tools (Fig.**2**A). The resulting datasets cover a broad range of common target and disease types (Fig.**3**C), providing a robust framework for systematic evaluation.

#### Construction of TRQA-lit

The TRQA-lit dataset specifically focuses on research findings related to therapeutic targets, aiming to build a question-answering (QA) dataset from literature corpora that summarizes the latest research progress for well-recognized therapeutic targets. We constructed the TRQA-lit dataset through an integrated approach combining LLM tools and human expertise via the following steps (Fig.**2**A, left):

1. A list of therapeutic targets was obtained based on the PubMed score from Pharos database[36] to ensure they represent research hotspots with accessible literature.
2. Human experts selected 90 targets and identified a comprehensive review paper for each, detailing its therapeutic potential and investigational progress.
3. Human experts then used LLM tools with self-defined prompts to extract question-answer pairs from these review papers, employing a unified question template focused on key considerations for target surveys.
4. To filter out trivial questions, a baseline model (Doubao without search) was used to answer the initial QA set, and correctly answered questions were removed.
5. The remaining questions underwent manual inspection and filtering by human experts for quality. For multi-choice questions where LLMs occasionally generated mislabeled options, Deepseek (DeepSeek-V3-250324) and Doubao-thinking (Doubao-1-5-Thinking-Pro-250415) were used to double-check answers against the literature corpus; inconsistent answers were then human-corrected.

As a result, TRQA-lit contains 172 multi-choice QAs (forming a core set for quick evaluation of models and humans) and 1,108 short-answer QAs, covering fundamental biology, disease biology, clinical medicine, and pharmacology (Fig.**3**A, left).

#### Construction of TRQA-db

While the scientific literature provides the main scientific discoveries related to therapeutic target research, the progress made in the industries (i.e., the competitive landscape) is also an important consideration when target-based discovery is being initiated. Therefore, we created the TRQA-db set from commercial databases, for systematically evaluating the effectiveness of information retrieval, integration, and reasoning among existing methods when addressing the competitive landscape investigation problem. To be specific, TRQA-db was constructed through the following steps (Fig.**2**A, right):

1. We obtained the drug discovery and development, patents, and clinical trial-related data for therapeutic targets in several emerging disease areas (i.e., schizophrenia, Crohn’s disease and non-alcoholic steatohepatitis), from the NextPharma database of PharmaCube ^1^ and the Synapse database of PatSnap ^2^.
2. We then wrote a list of questions covering key aspects of competitive landscape information.
3. Finally, the ground-truth answers were automatically extracted from the databases using human-defined rules.

TRQA-db contains 641 short-answer QAs, which mainly focus on key competitive information of drug R&D pipelines and clinical trials (Fig.**3**A, right).

#### Evaluation Methods

To robustly assess model performance across different question types, distinct evaluation methodologies were employed for multiple-choice and short-answer questions.

For multiple-choice questions, a two-step evaluation process ensured accurate assessment of option selection:

1. Option Extraction: The LLM was provided with the question, all available options, and the model’s generated answer. The LLM was then instructed to extract the option(s) from the provided choices that best matched its own generated answer. This mitigates issues where a model might generate a correct free-text answer that doesn’t perfectly match pre-defined option phrasing.
2. Correctness Validation: The extracted matched options were then compared against the golden (correct) answer to determine correctness based on an exact match.

This method allows for a flexible yet rigorous evaluation of how well the model identifies and selects the correct response among given choices.

For short-answer questions, a keyword-based recall evaluation was adopted to measure response completeness:

1. Keyword Extraction: An LLM was first utilized to identify and extract key concepts or keywords present in the golden (correct) answer. This step ensures that the evaluation focuses on the most critical pieces of information required for a complete answer.
2. Keyword Presence Check: For each extracted keyword, the LLM was then tasked with determining whether or not that specific keyword was mentioned or implied in the model’s generated output. This step allows for a granular assessment of which essential pieces of information are present in the response.
3. Keyword Recall Calculation: The keyword recall score was calculated by dividing the number of keywords present in the model’s output by the total number of keywords extracted from the golden answer, quantifying how comprehensively the model addressed the core information.

This methodology emphasizes providing relevant and comprehensive information over strict phrasing adherence, crucial for open-ended questions

Specific metrics were then applied to these evaluated responses. For TRQA-lit multiple-choice questions, accuracy is the main evaluation metric. For short-answer questions in both TRQA-lit and TRQA-db, models generate free-form responses; GPT-4o-mini is then used to extract keywords from these responses for comparison against keywords from ground-truth answers. Recall scores are primarily used, as source data (literature and databases) may not always contain the most cutting-edge progress, but are generally accurate.

#### Baselines

To evaluate OriGene’s performance against existing methods on TRQA and other public benchmark datasets, a comprehensive set of baseline models was selected, categorized into three groups:

- LLMs: We included DeepSeek-R1[29], DeepSeek-V3[19], GPT-41[23], o3-mini[37], Claude-3.7-Sonnet[3], and Gemini[17] as representative state-of-the-art LLMs, offering strong general knowledge and reasoning capabilities. DeepSeek-R1 and DeepSeek-V3 excel in advanced language understanding and reasoning tasks. Claude-3.7-Sonnet is a hybrid reasoning model, capable of quick responses or extended step-by-step thinking. GPT-41 provides advanced multimodal processing, while o3-mini is a time- and resource-efficient option. Gemini is known for its efficiency in complex tasks.
- General multi-agent: GPT-4o-search was selected as a general multi-agent baseline. This model integrates search capabilities with LLM reasoning, enabling it to dynamically retrieve and process external information.
- Biomedical multi-agent: Tx-Agent and Biomni(in core set) were chosen as a domain-specific baseline, designed explicitly for biomedical tasks. Its architecture is tailored for biological data analysis and target identification, providing a direct comparison to OriGene’s performance in this domain.
- Human Group: To establish a human baseline, we recruited a panel of nine experts in disease biology, chemical biology, or clinical medicine. The experts answered 172 multiple-choice questions from the TRQA-lit dataset, a subset selected for its manageable size. We organized the experts into three groups based on experience: senior doctoral candidates, recent doctoral graduates, and doctoral graduates with extensive work experience (Fig.**2**C). To generate three independent replicates of the baseline, the full set of questions was divided equally among the members within each group. Participants were permitted to use Google for reference but prohibited from using LLM or agent-based tools, with an instructed time of approximately two minutes per question.

### 4.5. Experimental Validation of Discovered Targets

#### Cell Lines and Culture Conditions

Human hepatocellular carcinoma (HCC) cell line Huh-7 and human colorectal cancer (CRC) cell line HCT116 were used. Cells were maintained in media (RPMI-1640) supplemented with 10% fetal bovine serum (FBS) and 1% penicillin-streptomycin, and cultured in a humidified incubator at 37°C with 5% CO2. Cell lines were routinely tested for mycoplasma contamination.

#### Peripheral Blood Mononuclear Cell (PBMC) Isolation and T Cell Culture

PBMCs were isolated from fresh human blood from Zhongshan Hospital Fudan University, with written consent and approval from the Institutional Review Board-approved protocols (B2021-381R and B2023-350R). PBMCs were isolated using Ficoll-Paque Plus (GE Healthcare) density gradient centrifugation. T-lymphocytes were subsequently isolated from PBMCs by using CD3 magnetic beads (Miltenyi Biotec) according to the manufacturer’s instructions.

#### Cell Viability and Proliferation Assays

Huh-7 (for GPR160 inhibitor) and HCT116 (for ARG2 inhibitor) cells were seeded in 96-well plates. After cell attachment, they were treated with various concentrations of GPR160 inhibitor or ARG2 inhibitor for a specified period. Cell viability was assessed by Propidium iodide using flow cytometry. Dose-response curves were generated, and half-maximal inhibitory concentrations (IC50) were calculated using non-linear regression analysis using R.

#### Patient-Derived Organoid (PDO) and Patient-Derived Tumor Fragments (PDTF)

Patient-derived tumor tissues (HCC and metastatic colorectal adenocarcinoma) were from Zhongshan Hospital Fudan University, with written consent and approval from the Institutional Review Board-approved protocols (B2021-381R and B2023-350R). Briefly, fresh tumor tissues were minced, digested, and the resulting cell clusters were embedded in Matrigel (Corning) droplets in 24- or 96-well plates. For drug testing, established PDOs were treated with GPR160 inhibitor (for HCC PDOs) or ARG2 inhibitor for 72h. Organoid viability was assessed using Propidium iodide.

#### Cancer Cell and T Cell Co-culture System

For GPR160 studies, Huh-7 HCC cells were cocultured with isolated human T-lymphocytes. Co-cultures were treated with GPR160 inhibitor or vehicle control. After 72h of treatment, T cells were harvested for analysis.

#### Flow Cytometry for T Cell Activation

T cells from co-culture experiments (with Huh-7 cells or HCC PDOs) were stained for surface markers. Data were acquired on a flow cytometer (BD LSRFortessa) and analyzed using FlowJo software (BD). Gating strategies were employed to identify T cell populations (CD3+) and assess the percentage of CD69+ cells within these populations.

#### Humanized Mouse Model

We strictly adhered to animal care principles and ethics and received approval from the Institutional Animal Care and Use Committee of the Shanghai Model Organisms Center (approval number 2019-0011). Immunodeficient mice (6 weeks old) were reconstituted with human PBMCs (1 × 10^7^ cells per mouse). After 7 days of humanization, mice were subcutaneously inoculated with Huh-7 cells (e.g., 5 × 10^6^ cells in 100 µL PBS/Matrigel mixture) into the flank. Mice were randomized into treatment groups (n=5) and treated with either GPR160 inhibitor (10µmol/kg) or vehicle control. Tumor volume was calculated using the formula: (Length × Width^2^)/2.

#### Statistical Analysis

Statistical significance between groups was determined using tests such as Student’s t-test or ANOVA using R, with p-values *<* 0.05 considered statistically significant. Specific tests and p-values are indicated in the figure legends or results section. Kaplan-Meier analysis with log-rank test was used for survival data.

## Acknowledgments

We thank our teammates Meihui Song, Wen Xiong, Jiabing Wang, and Haixiang Pei for their detailed guidance and technical feedback on the manuscript. We gratefully acknowledge the experts who contributed to benchmark question design, problem-solving and scoring: Jiaqiang Ma, Haichao Zhao, Guohe Song, Youpei Lin, Tiancheng Zhang, Hao Xiao, Xulang Peng, Rongyuan Wei, Jing Xing, Yuru Han, Sulin Zhang, Yiming Wen and Xinyue Shao. We also thank Wenhao Wang, Yueyang Tang, Jiahao Xu and Zhongmin Li for their contribution on MCP tools. Jiahao Tan, Xinyi Ma and Jiayue Cao for their contribution additional figure preparation, Mengyan Chen and Yanjie Huang for their valuable effort on database construction and Yinfu Wei for his contributions to platform development. This study has been supported by the Langang Laboratory [LGL-8888], the National Natural Science Foundation of China [62041209 to S.Z., 82441049 to Q.G., 823B2062 to Y.W.], Shanghai Sailing Program (22YF1460800) to D.W., the Young Elite Scientists Sponsorship Program by CAST [2023QNRC001 to S.Z.], the Science and Technology Commission of Shanghai Municipality [24510714300 to S.Z.], a locally commissioned task from the Shanghai Municipal Government to B.Z., Major Project of Guangzhou National Laboratory (grant no. GZNL2024A01015) to L.T. and F.T., S. Z. acknowledges funding from the Asian Young Scientist Fellowship. S.S. acknowledges funding from the Shanghai Artificial Intelligence Laboratory.

## Author Contributions

S.Z. led and supervised the project. S.Z., S.S., and Q.G. conceived the project. Z.Zhang, Z.Q., S.L., D.W., Z.Zhou, and Y.Wu coordinated development, built the agent architecture, and drafted major manuscript sections. Z.Q., Y.C., Z.Zhou, Z.Zhang, and Z.Zhong developed and benchmarked the multi-agent and LLM pipelines. Y.Wu and J.L. performed biological experiments and analyses. Y.L., K.W., and S.Song led the development of the biological tool modules and MCP services. X.W., Z.Shi, and K.W. further contributed to MCP tool development, contributed substantially to the analytical design and interpretation of the study. X.W., W.L. and Z.Zhong participated in manuscript revision. S.L., Y.Wang, W.Z., Q.L., B.D., and Y.He assembled the benchmark, performed analyses, prepared figures, and revised the manuscript with D.W. L.T. and F.T. provided constructive feedback on the framework, benchmark study design, and prebuilt workflow template design. Y.W., D.A., Z.Zhang, Y.L., T.Liu, and J.G. contributed to the design and deployment of the OriGene web product, including system architecture and interface development. J.C. further revised the manuscript and provided biological insights. Y.H., Z.Wang, Z.Wei, and R.M. extended the OpenTargets and MCP interfaces. Y.L. and C.O. generated illustrations. B.Z. and Y.Hu implemented test-time scaling and self-evolving algorithms. Q.F. maintained the server infrastructure, and D.A. managed project administration and platform implementation. S.Z., S.S., Z.Zhou, Z.Zhang, Z.Q., D.W., W.Y., S.L., J.Z., H.X., Y.C., J.C., and Y.Wu wrote the manuscript with input from all authors.

## Data and Materials Availability

The code will be released publicly on GitHub at https://github.com/GENTEL-lab/OriGene. The dataset will be available at https://huggingface.co/datasets/GENTEL-Lab/TRQA.

## Footnotes

1 https://www.pharmcube.com/en/product/nextpharma/index

2 https://synapse.patsnap.com/

## References

[1] J. Abramson, J. Adler, J. Dunger, R. Evans, T. Green, A. Pritzel, O. Ronneberger, L. Willmore, A. J. Ballard, J. Bambrick, et al. Accurate structure prediction of biomolecular interactions with alphafold 3. Nature, 630(8016):493–500, 2024.

[2] S. Aibar, C.B. González-Blas, T. Moerman, V. A. Huynh-Thu, H. Imrichova, G. Hulselmans, F. Rambow, J.-C. Marine, P. Geurts, J. Aerts, et al. Scenic: single-cell regulatory network inference and clustering. Nature methods, 14(11):1083–1086, 2017.

[3] Anthropic. Claude 3.7 sonnet system card. https://assets.anthropic.com/m/785e231869ea8b3b/original/claude-3-7-sonnet-system-card.pdf, February 2025. Accessed: 2025-06-01.

[4] M. Ashburner, C. A. Ball, J. A. Blake, D. Botstein, H. Butler, J. M. Cherry, A. P. Davis, K. Dolinski, S. S. Dwight, J. T. Eppig, et al. Gene ontology: tool for the unification of biology. Nature genetics, 25(1):25–29, 2000.

[5] J. Baek, S. K. Jauhar, S. Cucerzan, and S. J. Hwang. Researchagent: Iterative research idea generation over scientific literature with large language models. arXiv preprint 2404.07738, 2024.

[6] T. Barrett, S. E. Wilhite, P. Ledoux, C. Evangelista, I. F. Kim, M. Tomashevsky, K. A. Marshall, K. H. Phillippy, P. M. Sherman, M. Holko, et al. Ncbi geo: archive for functional genomics data sets—update. Nucleic acids research, 41(D1):D991–D995, 2012.

[7] S. K. Burley, H. M. Berman, G. J. Kleywegt, J. L. Markley, H. Nakamura, and S. Velankar. Protein data bank (pdb): the single global macromolecular structure archive. Protein crystallography: methods and protocols, pages 627–641, 2017.

[8] D. Carvalho-Silva, A. Pierleoni, M. Pignatelli, C. Ong, L. Fumis, N. Karamanis, M. Carmona, A. Faulconbridge, A. Hercules, E. McAuley, et al. Open targets platform: new developments and updates two years on. Nucleic acids research, 47(D1):D1056–D1065, 2019.

[9] S. W. Choi and P.F. O’Reilly. Prsice-2: Polygenic risk score software for biobank-scale data. Gigascience, 8(7):giz082, 2019.

[10] U. Consortium. Uniprot: a worldwide hub of protein knowledge. Nucleic acids research, 47 (D1):D506–D515, 2019.

[11] S. A. Dugger, A. Platt, and D. B. Goldstein. Drug development in the era of precision medicine. Nature reviews Drug discovery, 17(3):183–196, 2018.

[12] B. Gao, Y. Huang, Y. Liu, W. Xie, W.-Y. Ma, Y.-Q. Zhang, and Y. Lan. Pharmagents: Building a virtual pharma with large language model agents. arXiv preprint 2503.22164, 2025.

[13] S. Gao, R. Zhu, Z. Kong, A. Noori, X. Su, C. Ginder, T. Tsiligkaridis, and M. Zitnik. Txagent: An ai agent for therapeutic reasoning across a universe of tools. arXiv preprint 2503.10970, 2025.

[14] S. Gao et al. Empowering biomedical discovery with ai agents. arXiv preprint 2404.02831, July 2024. URL http://arxiv.org/abs/2404.02831. Accessed: Aug. 27, 2024.

[15] I. Gashaw, P. Ellinghaus, A. Sommer, and K. Asadullah. What makes a good drug target? Drug discovery today, 16(23-24):1037–1043, 2011.

[16] A. E. Ghareeb, B. Chang, L. Mitchener, A. Yiu, C. J. Szostkiewicz, J. M. Laurent, M. T. Razzak, A. D. White, M. M. Hinks, and S. G. Rodriques. Robin: A multi-agent system for automating scientific discovery. arXiv preprint 2505.13400, 2025.

[17] Google DeepMind. Gemini 2.5 pro. https://deepmind.google/technologies/gemini/pro/, May 2025. Accessed: 2025-06-01.

[18] J. Gottweis, W.-H. Weng, A. Daryin, T. Tu, A. Palepu, P. Sirkovic, A. Myaskovsky, F. Weis-senberger, K. Rong, R. Tanno, et al. Towards an ai co-scientist. arXiv preprint 2502.18864, 2025.

[19] D. Guo, D. Yang, H. Zhang, J. Song, R. Zhang, R. Xu, Q. Zhu, S. Ma, P. Wang, X. Bi, et al. Deepseek-r1: Incentivizing reasoning capability in llms via reinforcement learning. arXiv preprint 2501.12948, 2025.

[20] W. Helen. Bridging the translational gap: collaborative drug development and dispelling the stigma of commercialization. Drug discovery today, 21(2):299–305, 2016.

[21] K. Huang, Y. Jin, R. Li, M. Y. Li, E. Candés, and J. Leskovec. Automated hypothesis validation with agentic sequential falsifications. arXiv preprint 2502.09858, 2025.

[22] K. Huang, S. Zhang, H. Wang, Y. Qu, Y. Lu, Y. Roohani, R. Li, L. Qiu, G. Li, J. Zhang, et al. Biomni: A general-purpose biomedical ai agent. biorxiv, pages 2025–05, 2025.

[23] A. Hurst, A. Lerer, A. P. Goucher, A. Perelman, A. Ramesh, A. Clark, A. Ostrow, A. Welihinda, A. Hayes, A. Radford, et al. Gpt-4o system card. arXiv preprint 2410.21276, 2024.

[24] D. M. Hyman, B. S. Taylor, and J. Baselga. Implementing genome-driven oncology. Cell, 168 (4):584–599, 2017.

[25] M. Kanehisa, M. Furumichi, M. Tanabe, Y. Sato, and K. Morishima. Kegg: new perspectives on genomes, pathways, diseases and drugs. Nucleic Acids Research, 45(D1):D353–D361, 2017. doi: 10.1093/nar/gkw1092.

[26] M. J. Landrum, J. M. Lee, M. Benson, G. Brown, C. Chao, S. Chitipiralla, B. Gu, J. Hart, D. Hoffman, J. Hoover, et al. Clinvar: public archive of interpretations of clinically relevant variants. Nucleic acids research, 44(D1):D862–D868, 2016.

[27] J. M. Laurent, J. D. Janizek, M. Ruzo, M. M. Hinks, M. J. Hammerling, S. Narayanan, M. Ponnapati, A. D. White, and S. G. Rodriques. Lab-bench: Measuring capabilities of language models for biology research. arXiv preprint 2407.10362, 2024.

[28] C. Li, Z. Tang, W. Zhang, Z. Ye, and F. Liu. Gepia2021: integrating multiple deconvolution-based analysis into gepia. Nucleic acids research, 49(W1):W242–W246, 2021.

[29] A. Liu, B. Feng, B. Xue, B. Wang, B. Wu, C. Lu, C. Zhao, C. Deng, C. Zhang, C. Ruan, et al. Deepseek-v3 technical report. arXiv preprint 2412.19437, 2024.

[30] S. Liu, Y. Lu, S. Chen, X. Hu, J. Zhao, Y. Lu, and Y. Zhao. Drugagent: Automating ai-aided drug discovery programming through llm multi-agent collaboration. arXiv preprint 2411.15692, 2024.

[31] C. Lu, C. Lu, R. T. Lange, J. Foerster, J. Clune, and D. Ha. The ai scientist: Towards fully automated open-ended scientific discovery. arXiv preprint 2408.06292, Aug. 2024. URL http://arxiv.org/abs/2408.06292. Accessed: Aug. 28, 2024.

[32] Q. Lu, G. P. Livi, S. Modha, K. Yusa, R. Macarrón, and D. J. Dow. Applications of crispr genome editing technology in drug target identification and validation. Expert opinion on drug discovery, 12(6):541–552, 2017.

[33] D. Mendez, A. Gaulton, A. P. Bento, J. Chambers, M. De Veij, E. Félix, M.P. Magariños J. F. Mosquera, P. Mutowo, M. Nowotka, M. Gordillo-Marañón, F. Hunter, L. Junco, G. Mugumbate, M. Rodriguez-Lopez, F. Atkinson, N. Bosc, C. J. Radoux, A. Segura-Cabrera, A. Hersey, and A. R. Leach. Chembl: towards direct deposition of bioassay data. Nucleic Acids Research, 47 (D1):D930–D940, 2019. doi: 10.1093/nar/gky1075.

[34] V. Naumov, D. Zagirova, S. Lin, Y. Xie, W. Gou, A. Urban, N. Tikhonova, K. Alawi, M. Durymanov, F. Galkin, et al. Dora ai scientist: Multi-agent virtual research team for scientific exploration discovery and automated report generation. bioRxiv, 2025.

[35] J. Navarro Gonzalez, A. S. Zweig, M. L. Speir, D. Schmelter, K. R. Rosenbloom, B. J. Raney, C. C. Powell, L. R. Nassar, N. D. Maulding, C. M. Lee, et al. The ucsc genome browser database: 2021 update. Nucleic acids research, 49(D1):D1046–D1057, 2021.

[36] D.-T. Nguyen, S. Mathias, C. Bologa, S. Brunak, N. Fernandez, A. Gaulton, A. Hersey, J. Holmes, L. J. Jensen, A. Karlsson, M. Liu, S. Mani, A. Menon, J. P. Overington, S. Patel, B. L. Roth, H.B. Schiöth V. Sharma, T. Sheils, L. A. Sklar, N. Southall, O. Ursu, D. Vidović, A. Waller, J. Yang, A. Jadhav, and T. I. Oprea. Pharos: Collating protein information to shed light on the druggable genome. Nucleic Acids Research, 45(D1):D995–D1002, 2017. doi: 10.1093/nar/gkw1072.

[37] OpenAI. o3 system card. https://openai.com/index/o3-o4-mini-system-card/, September 2024. Accessed: 2025-06-01.

[38] S. M. Paul, D. S. Mytelka, C. T. Dunwiddie, C. C. Persinger, B. H. Munos, S. R. Lindborg, and A. L. Schacht. How to improve r&d productivity: the pharmaceutical industry’s grand challenge. Nature reviews Drug discovery, 9(3):203–214, 2010.

[39] D. Rein, B. L. Hou, A. C. Stickland, J. Petty, R. Y. Pang, J. Dirani, J. Michael, and S. R. Bowman. GPQA: A graduate-level google-proof q&a benchmark. In First Conference on Language Modeling, 2024. URL https://openreview.net/forum?id=Ti67584b98.

[40] J. Saltz, R. Gupta, L. Hou, T. Kurc, P. Singh, V. Nguyen, D. Samaras, K. R. Shroyer, T. Zhao, R. Batiste, et al. Spatial organization and molecular correlation of tumor-infiltrating lymphocytes using deep learning on pathology images. Cell reports, 23(1):181–193, 2018.

[41] J. Shi and M. G. Walker. Gene set enrichment analysis (gsea) for interpreting gene expression profiles. Current Bioinformatics, 2(2):133–137, 2007.

[42] K. Swanson et al. The virtual lab: Ai agents design new sars-cov-2 nanobodies with experimental validation. bioRxiv, Nov. 2024. URL https://www.biorxiv.org/content/10.1101/2024.11.01.621423v1.

[43] K. Tomczak, P. Czerwińska, and M. Wiznerowicz. Review the cancer genome atlas (tcga): an immeasurable source of knowledge. Contemporary Oncology/Współczesna Onkologia, 2015(1):68–77, 2015.

[44] H. Wang, Y. He, P. P. Coelho, M. Bucci, A. Nazir, B. Chen, L. Trinh, S. Zhang, K. Huang, V. Chandrasekar, et al. Spatialagent: An autonomous ai agent for spatial biology. bioRxiv, pages 2025–04, 2025.

[45] Z. Wei, Y. Ektefaie, A. Zhou, D. Negatu, B. B. Aldridge, T. B. Dick, M. Skarlinski, A. White, S. G. Rodriques, S. Hosseiniporgham, et al. Fleming: An ai agent for antibiotic discovery in mycobacterium tuberculosis. bioRxiv, pages 2025–04, 2025.

[46] J. White. Pubmed 2.0. Medical reference services quarterly, 39(4):382–387, 2020.

[47] C. H. Wong, K. W. Siah, and A. W. Lo. Estimation of clinical trial success rates and related parameters. Biostatistics, 20(2):273–286, 2019.

